# Leveraging both individual-level genetic data and GWAS summary statistics increases polygenic prediction

**DOI:** 10.1101/2020.11.27.401141

**Authors:** Clara Albiñana, Jakob Grove, John J. McGrath, Esben Agerbo, Naomi R. Wray, Thomas Werge, Anders D. Børglum, Preben Bo Mortensen, Florian Privé, Bjarni J. Vilhjálmsson

## Abstract

The accuracy of polygenic risk scores (PRSs) to predict complex diseases increases with the training sample size. PRSs are generally derived based on summary statistics from large meta-analyses of multiple genome-wide association studies (GWAS). However, it is now common for researchers to have access to large individual-level data as well, such as the UK biobank data. To the best of our knowledge, it has not yet been explored how to best combine both types of data (summary statistics and individual-level data) to optimize polygenic prediction. The most widely used approach to combine data is the meta-analysis of GWAS summary statistics (Meta-GWAS), but we show that it does not always provide the most accurate PRS. Through simulations and using twelve real case-control and quantitative traits from both iPSYCH and UK Biobank along with external GWAS summary statistics, we compare Meta-GWAS with two alternative data-combining approaches, stacked clumping and thresholding (SCT) and Meta-PRS. We find that, when large individual-level data is available, the linear combination of PRSs (Meta-PRS) is both a simple alternative to Meta-GWAS and often more accurate.

## 1. Introduction

Polygenic risk scores (PRSs) are a powerful approach to summarize the individual genetic liability to develop a specific disease. They are particularly useful for complex traits and diseases, such as psychiatric disorders^1^, as these are often highly polygenic^2^. This is because PRSs aggregate the small risk contributions from thousands of variants into a single score, summarizing their overall risk contribution^3^. Broadly, the existing polygenic prediction methods differ in the type of data they use for training, i.e. individual-level genotypes/dosages or GWAS summary statistics. Today, GWAS summary statistics are widely available for a broad range of diseases and traits in public databases, e.g. the GWAS catalog contains more than 1,400 summary statistics^4^. For psychiatric disorders, the Psychiatric Genomics Consortium (PGC) provides GWAS summary statistics based on ever larger sample sizes, as a result of meta-analyzing the individual efforts of many research groups worldwide. Furthermore, many GWAS summary statistics-based PRS methods are broadly used: Clumping and Thresholding (C+T)^5–7^, LDpred^8^ or more recent methods^9–13^, and have proven successful to identify individuals with significant increased risk of complex diseases such as coronary artery disease^14^.

Interestingly, many of these *external* GWAS summary statistics-based PRS methods approximate the results of the *internal* individual-level data approaches, making some assumptions in the process (e.g. LDpred-inf^8^ and sBLUP^15^ approximate the genomic BLUP^16^, assuming that linkage disequilibrium (LD) patterns in the external data from which the GWAS summary statistics were derived can be captured using an LD reference). Furthermore, phenotype definition, genetic architecture and/or technical artifacts may affect the prediction accuracy of the derived PRSs^17,18^. Using methods that fit prediction effect sizes jointly on internal individual-level data for training PRSs makes some of these assumptions unnecessary, which can lead to improved prediction accuracy^8,19^ e.g. Privé *et al*. found that prediction of height using penalized linear regression provides more accurate PRSs compared to C+T (LD clumping an p-value thresholding) when trained on individual-level data^20^. Indeed, there exist a number of powerful alternatives for deriving PRSs using individual-level data^20–25^. Until recently, most individual-level datasets have been small, especially in comparison to sample sizes achieved in GWAS meta-analyses, but cheaper genotyping has led to the generation of large genetic datasets (e.g. iPSYCH for psychiatric disorders^26^ and UK Biobank for a multitude of complex traits^27^). Therefore researchers often have access to large individual-level genetic data as well as large published GWAS summary statistics. However, most PRS methods train on either of these data types separately but not directly on both (although many methods do require individual-level data for hyper-parameter optimization). SCT is the only exception that we are aware of, as it does train directly on both types of data^7^. By combining and leveraging data, we aim to increase the training sample size of PRSs and, ultimately, their prediction accuracy.

In the current paper, we explore and compare different approaches of combining internal individual-level data and external GWAS summary statistics for polygenic prediction. Currently, the most widespread approach is combining the data at the level of GWAS summary statistics by meta-analyzing the marginal effect estimates of different studies, prior to training the PRS (Meta-GWAS). We believe this approach is reasonable when the individual-level data is small, but may discards its potential for training when larger sample sizes are available. Alternatively, SCT^7^ generates a range of C+T PRSs from the external GWAS summary statistics over a grid of hyper-parameters (e.g. LD clumping parameters and p-value thresholds) and then stacks these PRSs by fitting a penalized regression model using individual-level data. This results in a more accurate PRS compared to C+T provided sufficient training data sample size. Based on weighted average PRSs^28,29^, we propose a model with two independently generated PRS (Meta-PRS): an internal PRS, derived from the individual-level data; and an external PRS, derived from the GWAS summary statistics; and train the weights using linear regression on a validation dataset. We derive the PRSs with methods that work well for highly polygenic traits — namely we use BOLT-LMM^30^ for deriving the internal PRS and LDpred^8^ for the external PRS. We compare the prediction accuracy of the three approaches presented above (Meta-GWAS, SCT and Meta-PRS) through simulations and application to real data of psychiatric disorders and other complex diseases and traits, using individual-level data from two large cohorts (iPSYCH and UK Biobank) as well as large GWAS summary statistics that excluded these cohorts. Finally, we provide guidelines for optimizing accuracies of PRS in different scenarios, i.e. different degrees of polygenicity and sample size ratios between GWAS summary statistics and individual-level data.

## 2. Methods

### 2.1. Approaches for combining internal and external data

We investigated the difference in prediction performance of PRSs that are trained using both external GWAS summary statistics and internal individual-level genetic data, but combined through three different approaches (Table 1). In the first approach (Meta-GWAS), the internal individual-level data was used to derive GWAS summary statistics that were subsequently meta-analyzed with the external GWAS summary statistics and finally used for deriving PRSs. For the second approach (SCT) we used the external summary statistics to derive a large number of C+T scores, and the individual-level data to fit a penalized regression to linearly combine these C+T scores. In the third approach (Meta-PRS), the individual-level data and GWAS summary statistics were used for deriving two independent PRSs. We obtained a weighted average of the two PRSs by fitting a linear regression model.

**Table 1:** Overview of the compared data-combining approaches and data utilization.

| Combining approach | Individual-level data | GWAS summary statistics | Combining strategy | Validation | Test |
| --- | --- | --- | --- | --- | --- |
| **Meta-GWAS** | GWAS | - | $PRS = \sum_{i=1}^M Z_i \cdot x_i$ $Z_i = \frac{\sqrt{N_{int}} \cdot z_{int} + \sqrt{N_{ext}} \cdot z_{ext}}{\sqrt{N_{int} + N_{ext}}}$ | Select PRS parameters | Assess PRS prediction accuracy |
| **SCT** | Penalized regression of C+T scores | Grid C+T scores | $PRS = \sum_{j=1}^k w_j \cdot PRS_j$ | - | |
| **Meta-PRS** | Derive $PRS_{int}$ | Derive $PRS_{ext}$ | $PRS = w_{int} \cdot PRS_{int} + w_{ext} \cdot PRS_{ext}$ | Select PRS parameters and linear regression effect sizes | |

$M$ : number of SNPs, $Z$ : SNP effect size, $x$ : SNP effect allele count, $N$ : effective sample size $N_{eff} = 4/(1/N_{ca} + 1/N_{co})$ , $int$ : internal data, $ext$ : external data, $k$ : number of PRSs in grid, $w$ : regression weights.

In the three approaches, the individual-level data was split in training, validation and test subsets following a 5-fold cross-validation scheme (4-0.5-0.5; 80% training, 10% validation, 10% testing). The selection criteria for all method parameters was the parameter maximizing prediction accuracy in terms of prediction *R*^2^ in the validation data. Consequently, we obtained 5 estimates of PRS prediction performance for each method in the test subset and reported the mean. The standard error of the mean prediction accuracy was estimated through 10K bootstrap replicates of this mean.

### 2.2. Computing PRSs

#### 2.2.1. Meta-GWAS

We obtained GWAS summary statistics for the individual-level data using linear regression implemented in the function big_univLinReg, from the R package bigstatsr^31^. We used sex, age, genotyping batch and the first 20 principal components (PCs) of the dataset as covariates in the GWAS. We performed a sample size-based meta-analysis with the external GWAS summary statistics using the software METAL^32^. We computed PRSs using LDpred v1.0.10^8^ (note that this version already implements some of the improvements made in LDpred2^33^), using the infinitesimal model and 7 priors assuming a proportion of causal variants (p = 1, 0.3, 0.1, 0.03, 0.01, 0.003, 0.001). We used a LD radius of 500 variants to compute the LD reference panel. We then selected the LDpred PRS with p maximizing the prediction *R*^2^ in the validation set. We also computed PRSs with LD-clumping and p-value thresholding (C+T), selecting the score from a set of C+T PRSs that maximized the prediction *R*^2^ in the validation set. The C+T PRSs were generated from a grid of parameters: LD pairwise correlation *r*^2^ values (0.01, 0.05, 0.1, 0.2, 0.5, 0.8, 0.95), base window sizes (50, 100, 200, 500) and 50 p-value thresholds (depending on max. and min. p-value in summary statistics, on a log-log scale)^7^. For LD clumping, the SNP p-values were used as a selection variable i.e. for a pair of correlated SNPs, the SNP with the lowest p-value was kept. A total of 1,400 C+T PRSs were derived for each chromosome.

#### 2.2.2. SCT

We computed C+T PRSs using the external GWAS summary statistics and the same grid of parameters as in section 2.2.1. The final PRS was computed using the function snp_grid_stacking from the R package bigsnpr^7^, which performs penalized logistic regression, with the 1400 x 22 C+T scores as predictors and phenotypes as outcomes in the training set.

#### 2.2.3. Meta-PRS

To obtain the Meta-PRS, we first computed two independent PRSs: *PRS*_*int*_ and *PRS*_*ext*_. For *PRS*_*int*_, we obtained per-SNP prediction betas with BOLT-LMM^25^ (using the flag –predBetas) and computed the PRS as 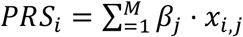, where *M* are the number of SNPs in the model, *β*_*j*_. For each sample and trait, we ran BOLT-LMM v2.3.4 using sex, age, genotyping batch and the first 20 PCs of the dataset as covariates. Depending on the polygenicity of the trait, BOLT-LMM computes a mixture-of-Gaussians prior on SNP effect sizes or the single-Gaussian BOLT-LMM-inf model, equivalent to best linear unbiased prediction (BLUP). The *PRS*_*ext*_ was computed with LDpred or C+T, as described in section 2.2.1. Finally, we defined the Meta-PRS with weights *w*_*ext*_ and *w*_*ext*_ as the linear combination of the two PRSs with these weights, as *MetaPRS* = *w*_0_ + *w*_*int*_*PRS*_*ext*_ + *w*_*int*_*PRS*_*ext*_ (lm function in R). To avoid overfitting, we trained the weights in a linear regression model in the validation data set (lm function in R). For the linear combination, we also used as weights the square root of the respective PRS training data sample size. In these cases, PRS were standardized prior to being combined. The latter use of weights is highlighted in the text, otherwise the weights in the Meta-PRS came from the linear regression model.

## 2.3. Data and quality control

### 2.3.1. iPSYCH data

We used genotype and phenotype data from the iPSYCH2012 case-cohort sample^26^. The iPSYCH2012 sample is nested within the entire Danish population born between 1981 and 2005, including 1,472,762 persons. Cases were identified as persons with schizophrenia (SCZ), autism (ASD), attention-deficit/hyperactivity disorder (ADHD), major depressive disorder (MDD) and anorexia nerviosa (AN); we identified controls as persons from the randomly selected cohort that were not diagnosed with any of the previous disorders. The genetics dataset consists of 78,050 individuals and 10,217,873 SNPs imputed following the RICOLPILI pipeline^34^. We computed KING-relatedness robust coefficient^35^ and excluded at random one of the individuals in the pairs > 3rd degree relatedness, resulting in 5,673 individuals excluded. We performed principal component analysis (PCA) following Privé, *et al*. 2020^36^ and obtained 30 PCs. We also identified 70,584 genetically homogeneous individuals based on these 30 PCs. We define homogeneous individuals as < 4.8 log(dist) units from the centre of the 30 PCs, calculated using the function dist_ogk from R package bigutilsr^36^. This resulted in a subset of 65,361 unrelated individuals of homogeneous ancestry. After removing SNPs with minor allele frequency (MAF) < 0.01 and Hardy–Weinberg p-value (*χ*^2^ (df = 1) test statistic pHWE) < 10^−6^, we restricted to the HapMap3 variants (https://www.sanger.ac.uk/resources/downloads/human/hapmap3.html). The final dataset was composed of 65,361 individuals and 1,184,138 SNPs.

### 2.3.2. UK Biobank data

We used genotype and phenotype data from the full release of the UK Biobank^27^, consisting of 488,377 individuals with genetic information. Specifically, we imported dosage data from BGEN files using the function snp_readBGEN from the R package bigsnpr^31^. We identified individuals with either self-reported or ICD-10 diagnosis for breast cancer (BC), coronary artery disease (CAD), type 2 diabetes (T2D) and major depressive disorder (MDD), setting the undiagnosed individuals as controls and restricting to women for breast cancer. We also identified individuals with standing height and body mass index (BMI) measurements to use as quantitative traits. We restricted the analysis to unrelated (as described in section 2.3.1) and “white British” genetic ancestry individuals. We removed SNPs with MAF < 0.01 and restricted to HapMap3 variants. The final dataset was composed of 337,475 individuals and 1,194,574 SNPs.

### 2.3.3. Simulations

We simulated case-control phenotypes using 1,194,574 HapMap3 SNPs and the subset of 337,475 unrelated European-ancestry individuals from the UK Biobank. The phenotypes were simulated with two different numbers of causal variants: *M*_*causal*_= 10k and 100k, representing polygenic traits. Each causal variant was assigned an effect size drawn from *N*(0, h^2^/*M*_*causal*_),where the heritability *h*^2^ = 0.5. The case-control status was assigned under a genetic liability model, with a simulated prevalence of 0.2. Each simulation scenario was repeated 5 times.

From the sample of individuals, 90% were used as the training set, 5% as the validation set and 5% as the test set. To represent scenarios with different sample sizes of the individual level data and GWAS summary statistics, the training set was further split randomly according to the following partitions: 10%-90%, 25%-75%, 50%-50%, 75%-25% and 90%-10%. One part was used to derive summary statistics and act as the external summary data, while the other part was used as individual-level data. The labels 9:1, 3:1, 1:1, 1:3, 1:9 used in the results reflect the sample size ratio of individual-level data (left) and GWAS summary statistics (right).

### 2.4. Prediction accuracy

The prediction accuracy of the PRSs was assessed in terms of squared correlation (*R*^2^) and area under the curve (AUC)^37^. The *R*^2^ was calculated for a model including the PRS and covariates (sex, age, genotyping batch and first 20 PCs) as explanatory variables and a model including only the covariates (without a PRS) as explanatory variables. The PRS prediction *R*^2^ was finally reported as 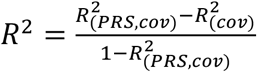 for the quantitative traits and transformed to the liability scale for the case-control data^38^. Additionally, the AUC was reported for the case-control data.

### 2.5. Code availability

The analysis pipeline was generated using gwf (https://docs.gwf.app/) and R scripts. All code used in this project is available at https://drive.google.com/drive/folders/1u6U55e8MERt3zzbQ3OQbiNJD5lLqGtUx?usp=sharing.

## 3. Results

### 3.1. Performance on simulated data

We evaluated the prediction accuracy of the PRSs using simulated data to explore the relationship between the combining approaches and the training sample size. Using the UK Biobank genetic data, we simulated traits with 10,000 (10k) and 100,000 (100k) causal SNPs, aiming at representing the polygenicity range of complex traits, and different sizes of training sample (10%, 25%, 50%, 75% and 90% of N ∼ 300,000 individuals) of individual-level data (internal) and GWAS summary statistics (external). First, we compared the prediction accuracy of PRSs trained only on internal data (using BOLT-LMM) or external data (using C+T or LDpred) in terms of mean prediction *R*^2^ (Fig. 1A) and AUC (Supplementary Fig. 1). For all simulated scenarios, the BOLT-LMM outperformed other methods, with a larger relative improvement in the simulations with 10k causal SNPs. The comparison between the GWAS summary statistics-based methods resulted in C+T being generally preferred in the simulations with 10k and LDpred in the ones with 100k causal SNPs. These results highlight the benefits of using the individual-level data for training PRSs over the derived GWAS summary statistics.

**Figure 1.**
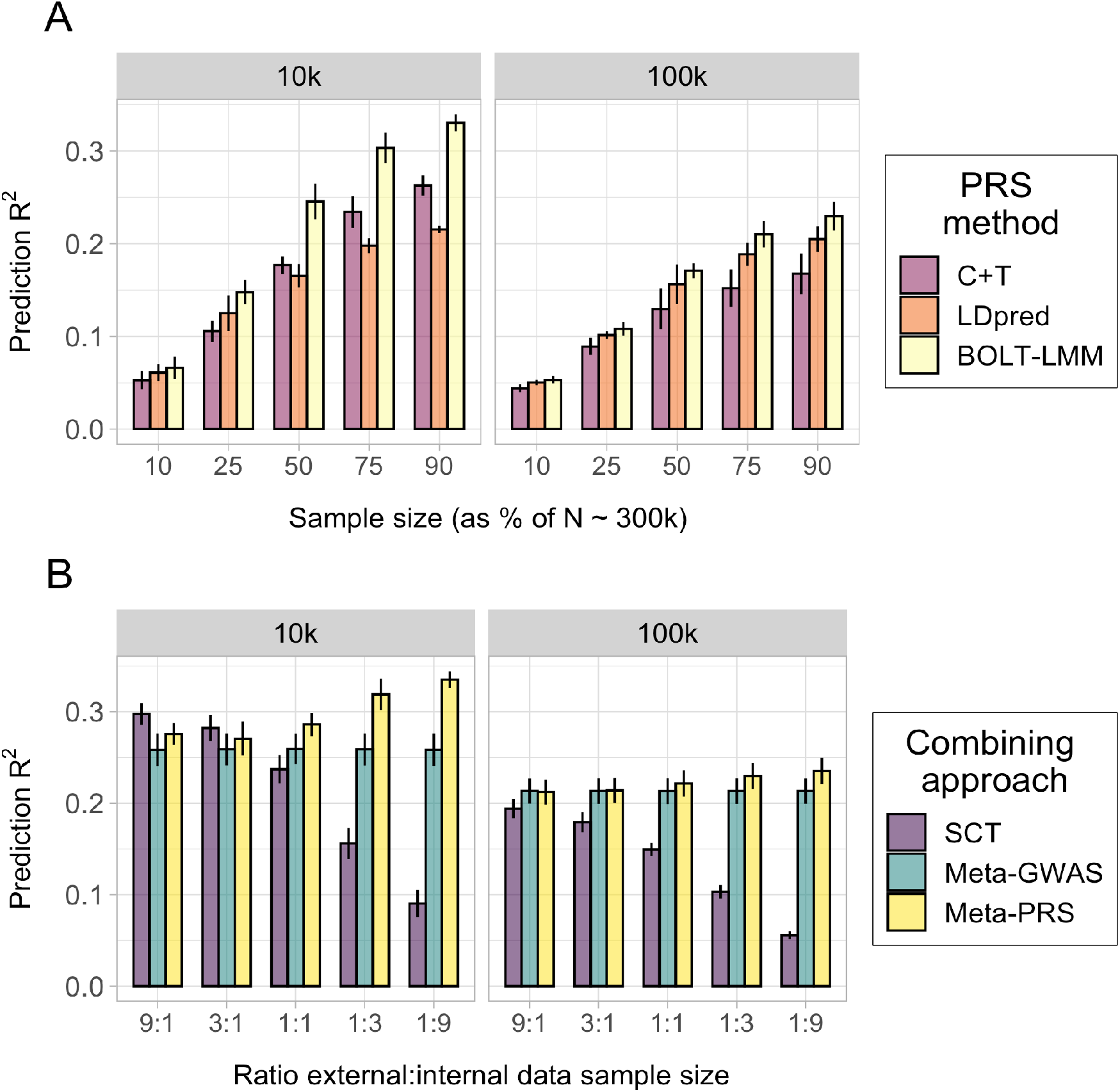
Prediction accuracy of the PRSs in the simulation study. Each panel displays the mean and 95% CI of the PRS prediction *R*^2^ (y-axis) for each data combining approach. The traits were simulated from a liability threshold model with 10,000 (10k) and 100,000 (100k) causal SNPs and heritability *h*^2^ of 0.5, and case-control status was inferred from a disease prevalence of 0.2. Mean and 95% CI of prediction *R*^2^ were obtained from 10k non-parametric bootstrap samples of 5 independent replicates. A) Effect of training sample size in the PRSs prediction accuracy. The x-axis indicates the percentage of individuals from the total training set (N = 303,728) used as individual-level data for BOLT-LMM or GWAS summary statistics for C+T and LDpred. B) Effect of the ratio between internal and external data in the combining approaches. The x-axis indicates the relative amount of external vs. internal data, e.g. 3:1 indicates a scenario where the external data was 25% and the internal data was 75% of the total sample. Fig. 1 is a simplified version of Supplementary Fig. 2, selecting a single method per combining approach between C+T and LDpred, where the method maximizing mean prediction *R*^2^ was selected.

We also compared the prediction accuracy of PRSs using different data-combining approaches (SCT, Meta-GWAS and Meta-PRS) in the simulated traits (Fig. 1B, Supplementary Fig. 2). The external and internal datasets were matched to create combinations with different ratios of each data type (9:1, 3:1, 1:1, 1:3, 1:9; e.g. 3:1 indicates a scenario where the external data was 75% and the internal data was 25% of the total N ∼ 300k individuals in the training set). For Meta-PRS, we observed a positive relation between the size of the internal data and the mean prediction *R*^2^. The opposite was observed for SCT, where larger external datasets provided larger mean predictions. The ratio of data showed no effect for Meta-GWAS, with constant prediction *R*^2^ along the simulated ratios (Fig. 1B). These results indicated that it was possible to optimize PRS prediction accuracy by selecting a data-combining approach depending on the sample size ratio between the available internal and external data. While the classical Meta-GWAS was a valid strategy in ratios of 1:1, scenarios with a more skewed ratio benefit from approaches like Meta-PRS and SCT, which use the individual-level data for training.

Aiming to simplify the construction of the Meta-PRS, we attempted to use the square root of the effective sample size 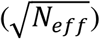 to weight the internal and external PRSs. This simplified version of Meta-PRS is faster and does not need of a validation dataset. In the previously-described simulated scenarios, we compared the mean prediction *R*^2^ of PRSs weighted by 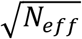 and PRSs weighted by linear regression effect sizes (using a validation dataset). We only observed a small increase in mean prediction *R*^2^ in the scenarios with large individual-level data (ratios 1:3 and 1:9), with the other remaining the same (Supplementary Fig. 3).

### 3.2. Performance on real data

We investigated the prediction accuracy of the data-combining approaches (Meta-PRS, SCT and Meta-GWAS) in real complex traits using internal individual-level data from large genotype cohorts (iPSYCH^26^ and the UK Biobank^27^) and external GWAS summary statistics without samples from these two cohorts. The set of traits selected included the six major psychiatric disorders (ASD, ADHD, MDD, BD, SCZ and AN), three other complex diseases (BC, T2D and CAD) and two continuous complex traits (height and BMI) (Table 2). The set of SNPs used for each trait was the intersection between the SNPs in the individual-level data, GWAS summary statistics and the 1,440,616 HapMap3 SNPs.

**Table 2:** Summary of real datasets. Sample sizes (cases / controls for binary traits) of the individual-level datasets for the 12 complex traits, along with the sample sizes of the corresponding GWAS summary statistics. The GWAS summary statistics selected did not include samples overlapping with the individual-level datasets used here. The table reflects sizes of European ancestry, unrelated samples (see Methods) and the ratios are based on effective sample sizes.

| Traits | Individual-level dataset | Individual-level sample size | GWAS sample size | Ratio | Overlapping SNPs |
| --- | --- | --- | --- | --- | --- |
| Attention deficit hyperactivity disorder (ADHD)<sup>39</sup> | iPSYCH | 17,072 / 25,982 | 4,225 / 11,012 | 3.4:1 | 1,105,731 |
| Autism spectrum disorder (ASD)<sup>40</sup> |  | 14,682 / 26,033 | 5,305 / 5,305 | 3.5:1 | 1,177,564 |
| Anorexia Nervosa (AN)<sup>41</sup> |  | 3,181 / 26,282 | 11,940 / 33,731 | 1:3.1 | 1,134,823 |
| Schizophrenia (SCZ)<sup>42</sup> |  | 2,701 / 26,277 | 21,169 / 28,117 | 1:4.9 | 1,183,697 |
| Bipolar Disorder (BD)<sup>43</sup> |  | 1,429 / 26,311 | 20,040 / 30,874 | 1:9 | 1,183,744 |
| Major depressive disorder (MDD)<sup>44</sup> |  | 22,469 / 25,882 | 229,897 / 544,204 | 1:13.4 | 1,094,603 |
| Height<sup>45</sup> | UK Biobank | 336,750 | 253,288 | 1.3:1 | 1,000,417 |
| Body mass index (BMI)<sup>46</sup> |  | 336,381 | 339,224 | 1:1 | 1,003,044 |
| Type 2 diabetes (T2D)<sup>47</sup> |  | 18,857 / 318,618 | 26,676 / 132,532 | 1:1.2 | 1,100,399 |
| Major depressive disorder (MDD)<sup>48</sup> |  | 28,626 / 308,849 | 45,396 / 97,250 | 1:1.2 | 1,091,232 |
| Coronary artery disease (CAD)<sup>49</sup> |  | 11,529 / 325,946 | 60,801 / 123,504 | 1:3.7 | 1,093,989 |
| Breast cancer (BC)<sup>50</sup> |  | 12,024 /<br>169,207 | 122,977 /<br>105,974 | 1:5.1 | 1,098,351 |

No single combining approach provided the largest mean prediction *R*^2^ for all traits (Fig. 2) or AUC (Supplementary Fig. 4) for all traits. In the cases where the sample size of individual-level data was larger than the summary statistics (int > ext), Meta-PRS increased mean prediction *R*^2^ over SCT and Meta-GWAS for height, while both Meta-GWAS and Meta-PRS had similar results for ASD and ADHD, with large and overlapping CIs. In the cases with equal data training sample sizes (1:1), Meta-PRS increased prediction accuracy over Meta-GWAS and SCT for BMI and T2D, while the results for Meta-GWAS and Meta-PRS were similar for MDD UKB. Finally, in the cases where the sample size of the GWAS summary statistics was larger than the individual-level data (ext > int) the results were also diverse. For AN, CAD, SCZ, BD and MDD iPSYCH there was no major difference between Meta-GWAS and Meta-PRS. However, for BC, the data-combining approach with the largest mean prediction *R*^2^ was SCT.

**Figure 2.**
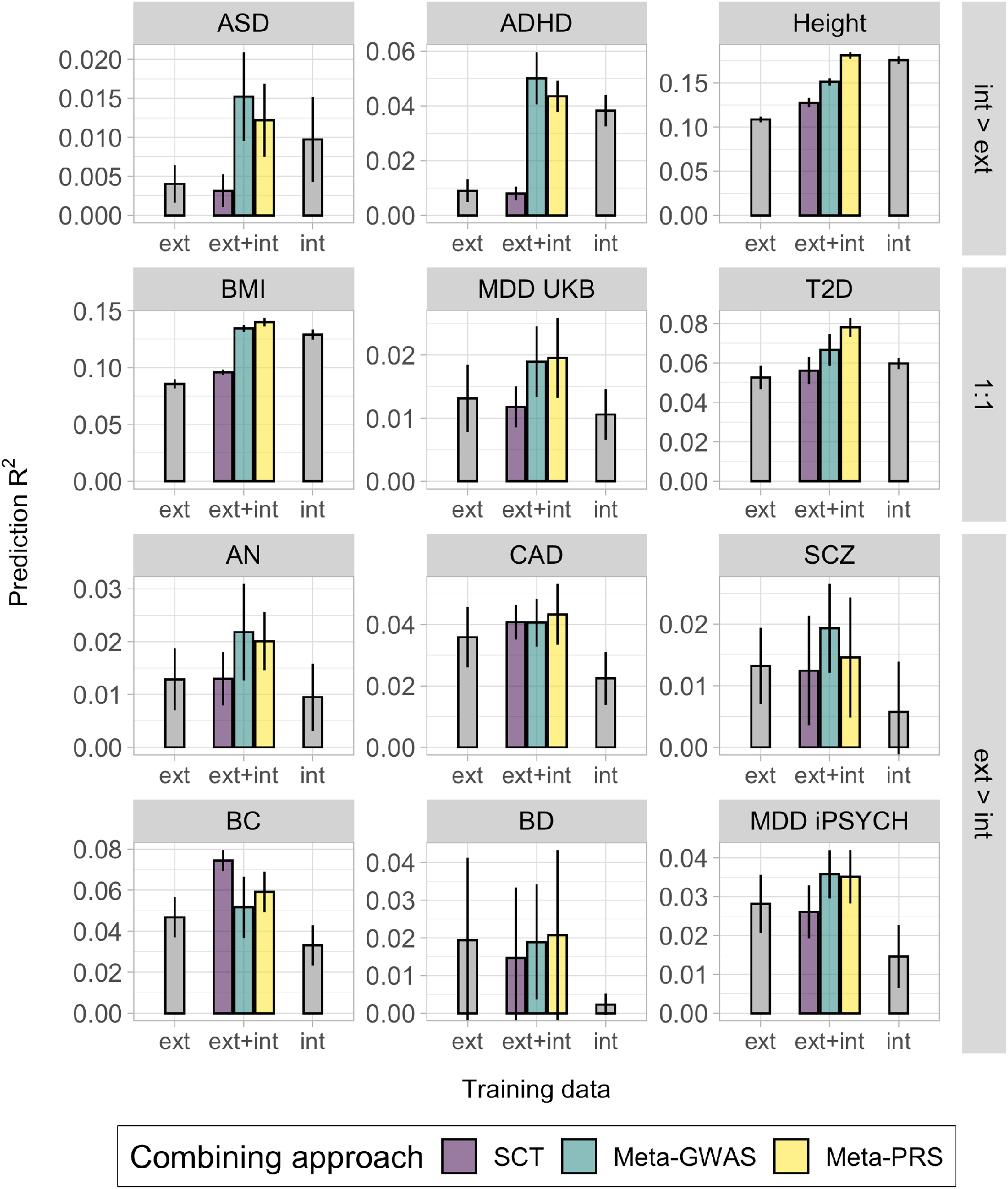
Prediction accuracy of the combining approaches in 12 complex traits from iPSYCH and UK Biobank. Each panel displays the mean and 95% CI of the PRS prediction *R*^*2*^ (y-axis) for each data combining approach, of PRS trained on individual-level data (int), GWAS summary statistics (ext) or both (ext+int) (x-axis). The prediction *R*^*2*^ was transformed to the liability-scale using a population prevalence of 0.01 (ASD), 0.05 (ADHD), 0.15 (MDD UK Biobank), 0.05 (T2D), 0.01 (AN), 0.03 (CAD), 0.01 (SCZ), 0.07 (BC), 0.01 (BD) and 0.08 (MDD iPSYCH). The methods noted as int and ext were fitted using BOLT-LMM with individual-level data and LDpred or C+T with GWAS summary statistics, respectively. For simplification, only the ext PRS with larger mean prediction *R*^*2*^ is shown, the full results are available in the Supplementary Fig.5. Mean and 95% CI of the prediction *R*^*2*^ were obtained from 10k non-parametric bootstrap samples of the 5 cross-validation subsets.

Generally, the Meta-GWAS showed a larger mean prediction *R*^2^ than Meta-PRS for the psychiatric disorders, though with large and overlapping CIs. This was independent of the sample size ratio of internal vs. external data. For most outcomes validated in the UK Biobank data, the most accurate approach was Meta-PRS, where the largest improvement was for height, BMI and T2D. For these outcomes the internal effective sample size was larger than for most of the other outcomes. BC was the only trait where SCT led to the most predictive PRS, even though the ratio internal:external was similar to other traits like CAD.

The PRS method-specific results showed a preference of LDpred over C+T in 6/12 traits, both in PRS trained on external or meta-analyzed summary statistics (Supplementary Fig. 5), while for the rest of the traits both methods had similar results. We also compared the Meta-PRS constructed with linear regression weights to the one weighed by effective sample sizes 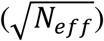 of training data (Supplementary Fig. 6). As in the simulations, we only observed an increase in mean prediction *R*^2^ in the traits with large individual-level data (height and BMI). In the rest of the traits, there was no preference for a specific weight type. The use of 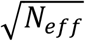 as weights is therefore recommended for these traits, as it does not require a validation set.

## 4. Discussion

With genetic data now available to researchers as both large individual-level datasets and GWAS summary statistics, we want to understand how to best combine these two types of data to optimize polygenic prediction. With this aim, we have evaluated the predictive performance of PRSs generated with different data-combining approaches: Meta-GWAS, SCT and Meta-PRS. We find that the simple approach of combining two different PRSs (Meta-PRS), trained on individual-level data and GWAS summary statistics separately, may yield more accurate PRSs than Meta-GWAS, particularly in the cases with sufficiently large individual-level datasets. We observe this in simulated data, where Meta-PRS consistently increases the mean prediction *R*^2^ over the widely used Meta-GWAS approach, and in the real complex traits with a large individual-level dataset e.g. height, BMI, and T2D. Another advantage of Meta-PRS is that it allows to combine multiple pre-calculated PRSs, irrespective of prediction method. When validation data is not available, we show that one can use the square root of the training sample sizes as weights. The same approach could also be used to combine multiple PRSs (e.g. in the PGS Catalog^51^), being standardized and averaged together with their corresponding training sample sizes. As an alternative approach, the scores in Meta-PRS could be weighted using MT-BLUP^52^.

In the case of BC, which has several large effects and relatively low polygenicity, the SCT PRS prediction is the most accurate, presumably because it relies more on variant thinning. For psychiatric disorders, we found that the Meta-GWAS generally yielded the most predictive PRSs, despite these disorders being very polygenic and often having relatively large individual-level data sample sizes. The results for the psychiatric disorders are contrary to what we expected based on our simulations regarding the preferred data-combining approach, although we note that the expected relative improvement of Meta-PRS over Meta-GWAS is small if polygenicity is large. Nevertheless, we want to highlight that leveraging the two types of data (individual-level data and GWAS summary statistics) always increased the prediction performance of PRSs over not combining data, even in the cases where either of these were small.

Our simulations represent an idealized scenario where we assume that the genetic architecture is invariant between cohorts/samples (i.e. genetic correlation is 1). Studies have shown that psychiatric disorders can be quite heterogenous between cohorts^18^, especially for the iPSYCH data where Schork *et al*. 2019^53^ estimated the genetic correlation for psychiatric disorders between external and iPSYCH samples to be between 0.5-0.8. Given that we found that Meta-GWAS provided more accurate predictions in the iPSYCH data, it may suggest that it is more robust to disease heterogeneity than Meta-PRS. However, if the genetic architecture is similar between samples (high genetic correlation), we expect Meta-PRS to have the advantage given even larger individual-level data sample sizes. Similar to disease heterogeneity, differences in genetic ancestry between the training and testing data can also decrease the prediction accuracy of PRSs^17^. In the case of ancestry heterogeneity, the linear combination of PRS trained independently on different ancestries improves prediction for admixed individuals^54^, but the extent to which these sample heterogeneities affect each of the prediction accuracy in the compared data-combining approaches should be further studied.

In Meta-PRS we combined the BOLT-LMM and LDpred (or C+T) predictions, and therefore the results may not be fully generalizable to other methods e.g. a more accurate method may lead to more accurate Meta-GWAS scores. Nevertheless, given that LDpred generally performs well for polygenic traits in independent comparisons^55,56^, we believe it acts as a good proxy for other similar methods, such as lasso regression^9^, SBayesR^11^, and PRS-CS^10^. In the case of individual-level data and low polygenicity, L1-penalized regression may also provide more accurate PRSs than BOLT-LMM^20^.

In summary, we found that a simple additive model of two polygenic scores (Meta-PRS) often outperformed the accuracy of approaches that first meta-analyzed SNP effects (Meta-GWAS) in highly polygenic traits. Fundamentally, the improvement in Meta-PRS prediction accuracy stems from the fact that methods that train a polygenic prediction model on individual-level data have access to more training information than methods that only train on a summary of this data and usually make fewer assumptions. However, Meta-GWAS has the advantage that each effect estimate is updated separately, possibly making it more robust to small sample sizes and changes in genetic architecture.

## Acknowledgments

This study was funded by grants from The Lundbeck Foundation (R102-A9118, R155-2014-1724, and R248-2017-2003) and The Danish National Research Foundation (Niels Bohr Professorship to Prof. John J. McGrath). The authors gratefully acknowledge the Psychiatric Genomics Consortium (PGC) and the research participants and employees of 23andMe, Inc. for providing the summary statistics. All of the computing for this project was performed on the GenomeDK cluster. We would like to thank GenomeDK and Aarhus University for providing computational resources and support that contributed to these research results. This research has been conducted using the UK Biobank Resource under Application Number 41181.

## Supplementary figures

**Supplementary Figure 1.**
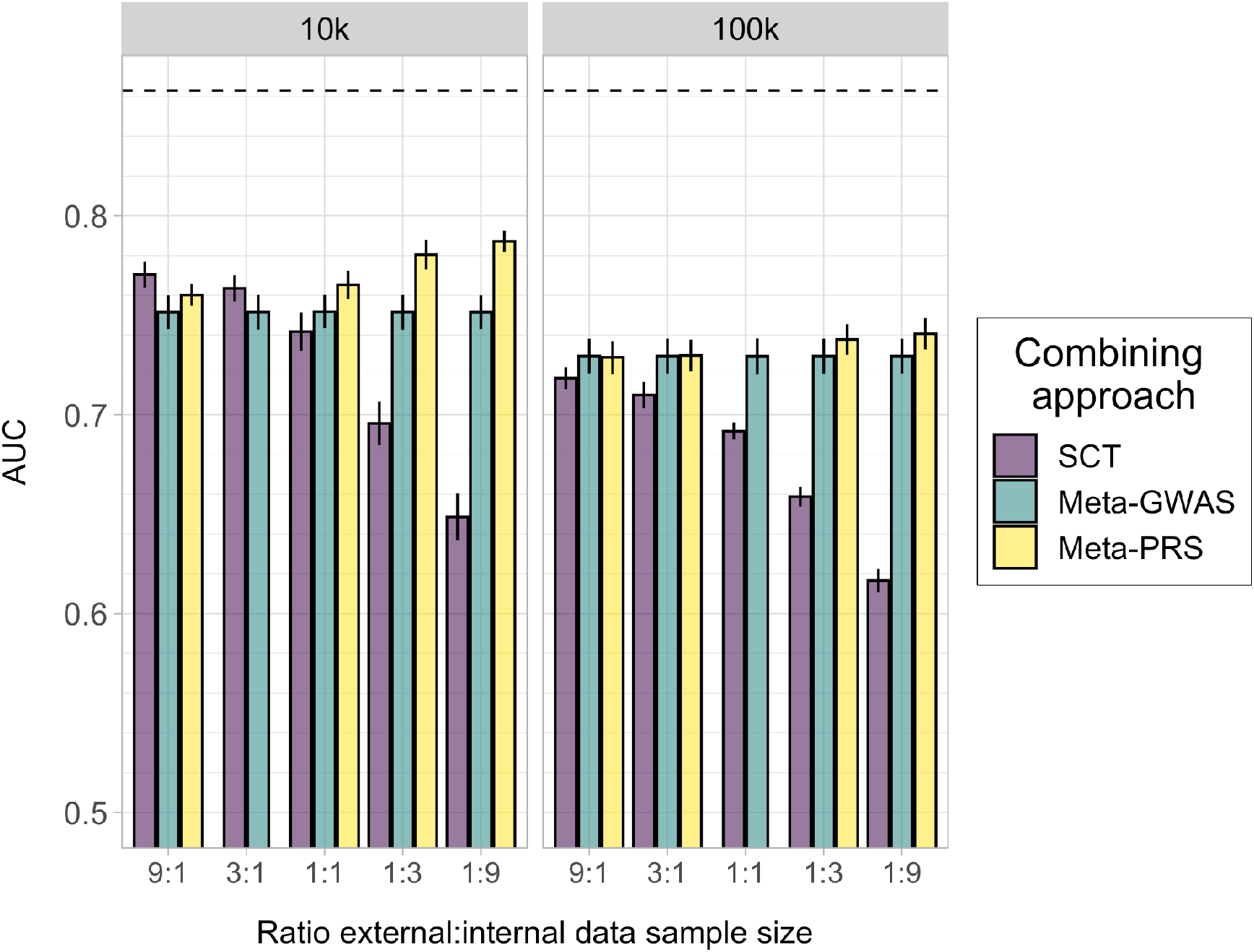
Prediction accuracy of the data-combining approaches in the simulated data in terms of AUC. Each panel displays the mean and 95% CI of the PRS AUC (y-axis) for each data-combining approach. The traits were simulated from a liability threshold model with 10,000 (10k) and 100,000 (100k) causal SNPs and heritability h^2^ of 0.5, and case-control status was inferred from a disease prevalence of 0.2. Mean and 95% CI of AUC were obtained from 10k non-parametric bootstrap samples of 5 independent replicates. The black line represents the AUC_max_(0.852) for these simulations^57^. The x-axis indicates the relative amount of external vs. internal data, e.g. 3:1 indicates a scenario where the external data was 25% and the internal data was 75% of the total sample (N = 303,728).

**Supplementary Figure 2.**
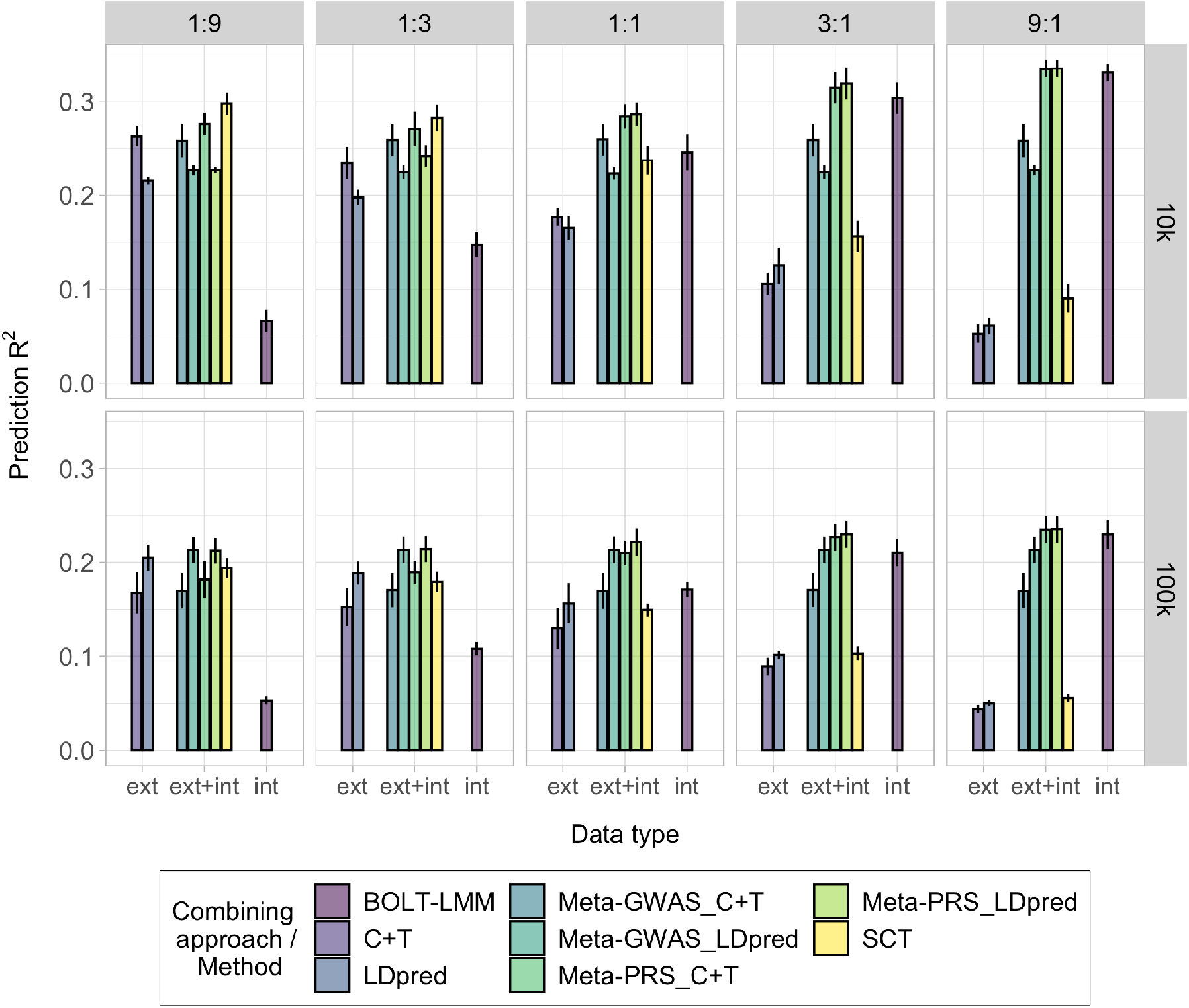
Prediction accuracy of the data-combining approaches using different GWAS summary statistics-based PRS methods in the simulated data. Each panel displays the mean and 95% CI of the PRS R^2^ (y-axis) for each data-combining approach and PRS method, of PRSs trained on individual-level data (int), GWAS summary statistics (ext) or both (ext+int) (x-axis). In the case of Meta-GWAS, C+T and LDpred were used on the meta-analyzed summary statistics and in Meta-PRS, C+T and LDpred were used to compute the external PRS. The traits were simulated from a liability threshold model with 10,000 (10k) and 100,000 (100k) causal SNPs and heritability h^2^ of 0.5, and case-control status was inferred from a disease prevalence of 0.2. Mean and 95% CI of prediction R^2^ were obtained from 10k non-parametric bootstrap samples of 5 independent replicates.

**Supplementary Figure 3.**
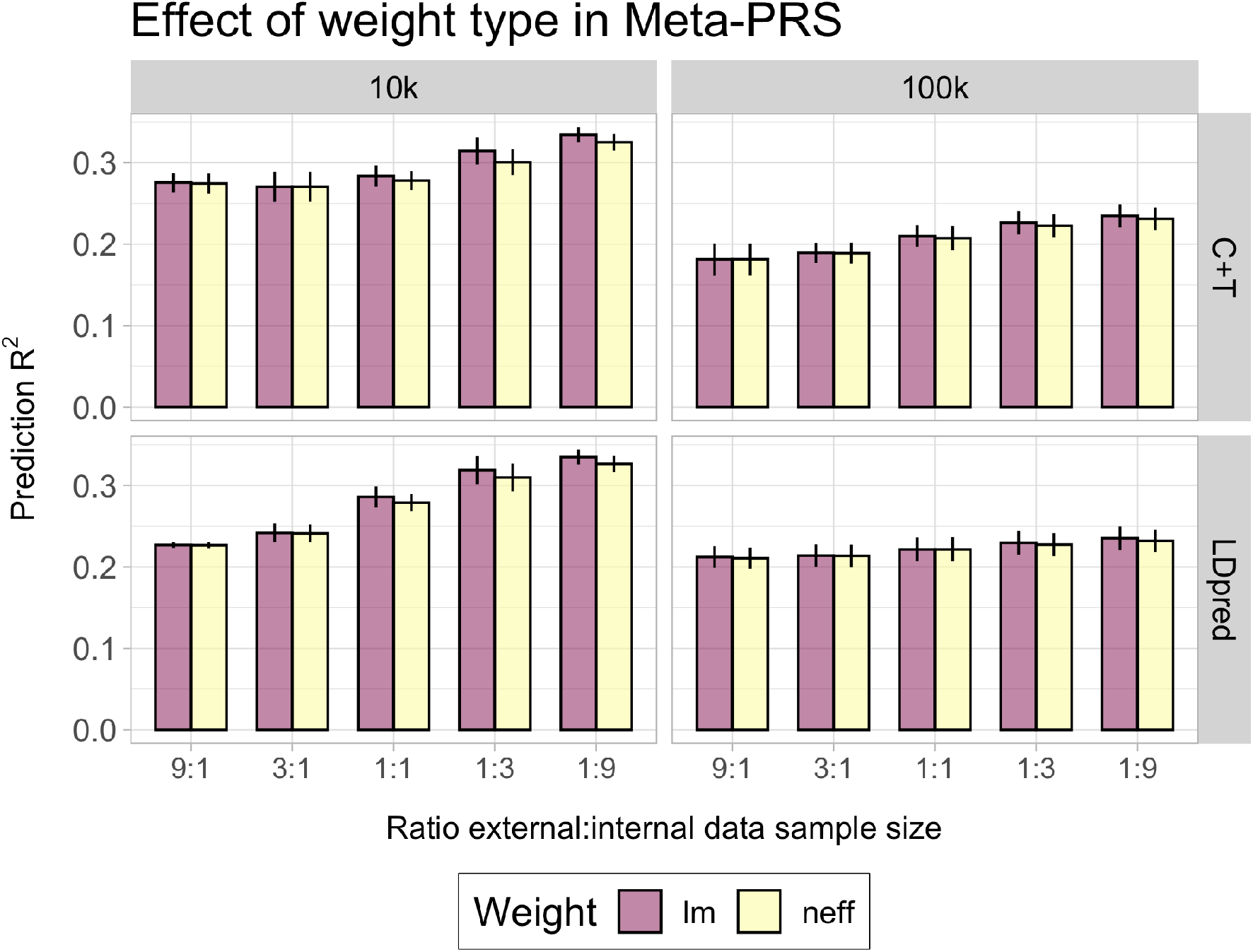
Prediction accuracy of Meta-PRS using different weight types in the simulated data. Each panel displays the mean and 95% CI of the PRS prediction R^2^ (y-axis) for Meta-PRS in each simulated scenario using either C+T or LDpred to generate the external PRS. The weights were obtained using linear regression (lm) or the square root of the training effective sample size (neff). In the case of the linear regression, the weights are trained in an independet validation dataset (see Table 1). The traits were simulated from a liability threshold model with 10,000 (10k) and 100,000 (100k) causal SNPs and heritability h^2^ of 0.5, and case-control status was inferred from a disease prevalence of 0.2. Mean and 95% CI of prediction R^2^ were obtained from 10k non-parametric bootstrap samples of 5 independent replicates.

**Supplementary Figure 4.**
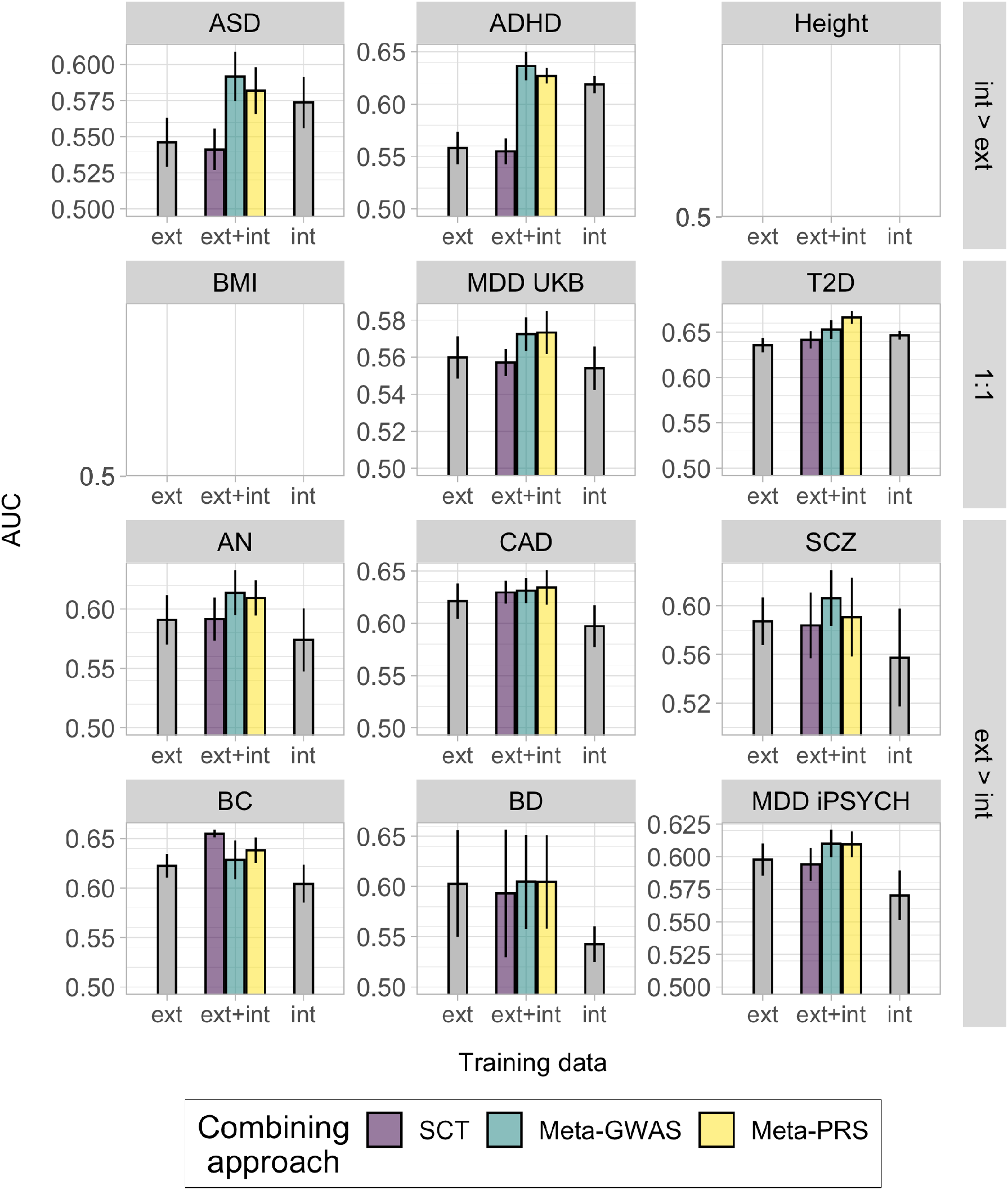
Prediction accuracy of the data-combining approaches in 12 complex traits from iPSYCH and UK Biobank. Each panel displays the mean and 95% CI of the PRS AUC (y-axis) for each data-combining approach, of PRS trained on individual-level data (int), GWAS summary statistics (ext) or both (ext+int) (x-axis). The methods noted as int and ext were fitted using BOLT-LMM with individual-level data and LDpred or C+T with GWAS summary statistics, respectively. For simplification, only the ext PRS with larger mean prediction R^2^ is shown. Mean and 95% CI of the AUC were obtained from 10k non-parametric bootstrap samples of the 5 cross-validation subsets.

**Supplementary Figure 5.**
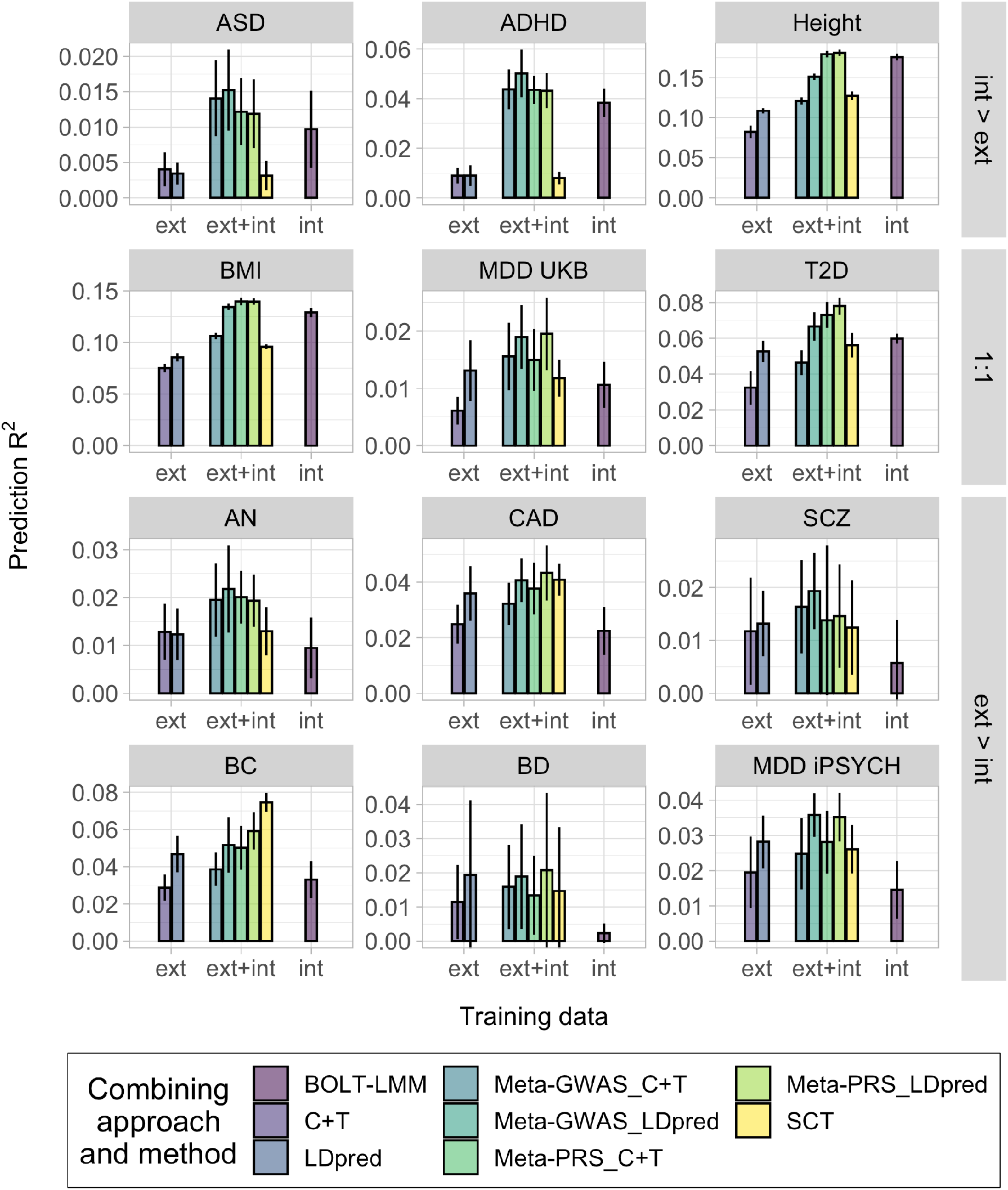
Prediction accuracy of the data-combining approaches using different GWAS summary statistics-based PRS method in 12 complex traits from iPSYCH and UK Biobank. Each panel displays the mean and 95% CI of the PRS R^2^ (y-axis) for each data-combining approach and PRS method, of PRSs trained on individual-level data (int), GWAS summary statistics (ext) or both (ext+int) (x-axis). In the case of Meta-GWAS, C+T and LDpred were used on the meta-analyzed summary statistics and in Meta-PRS, C+T and LDpred were used to compute the external PRS. The prediction R^2^ was transformed to the liability-scale using a population prevalence of 0.01 (ASD), 0.05 (ADHD), 0.15 (MDD UKB), 0.05 (T2D), 0.01 (AN), 0.03 (CAD), 0.01 (SCZ), 0.07 (BC), 0.01 (BD) and 0.08 (MDD iPSYCH). Mean and 95% CI of the AUC were obtained from 10k non-parametric bootstrap samples of the 5 cross-validation subsets.

**Supplementary Figure 6.**
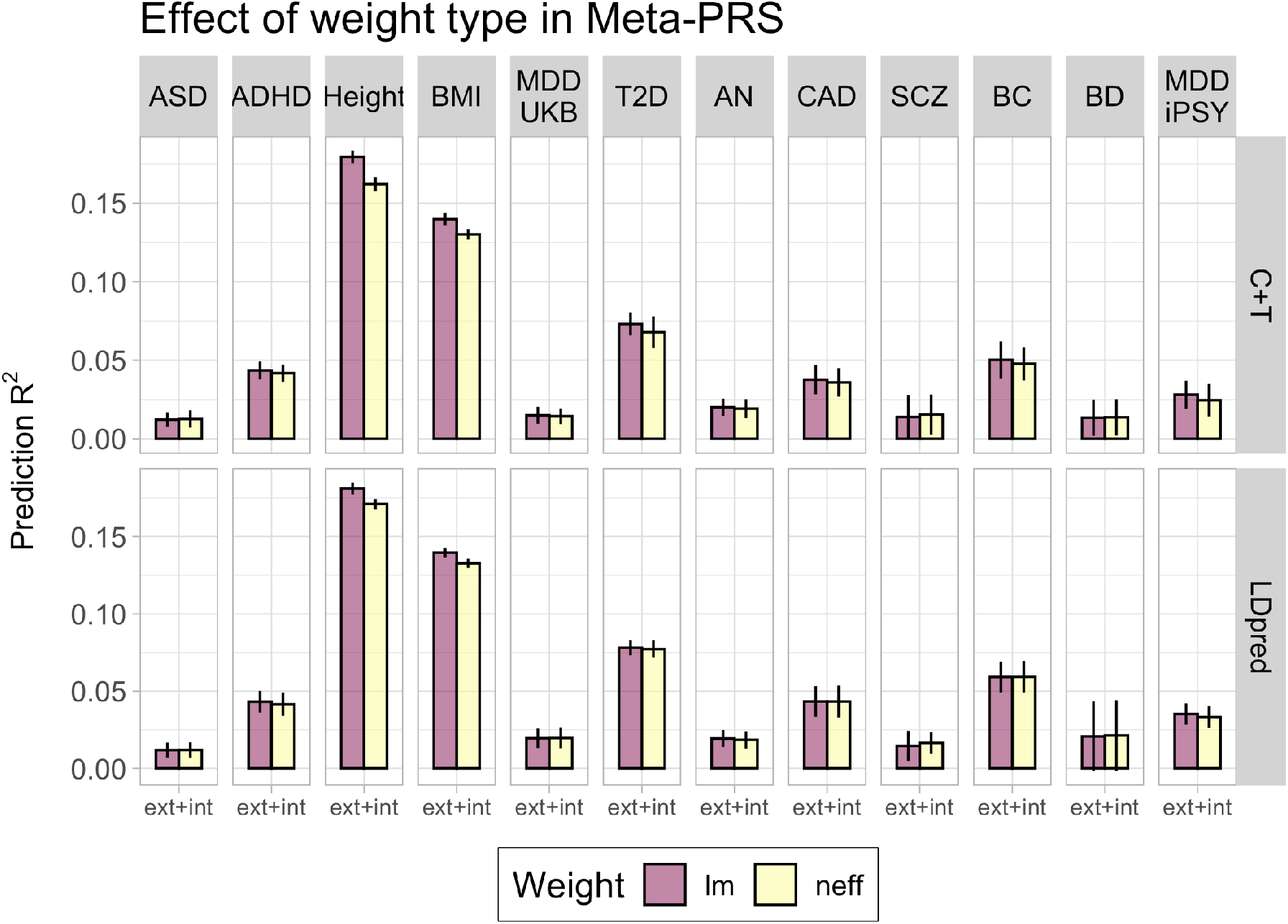
Prediction accuracy of Meta-PRS using different weight types in 12 complex traits from iPSYCH and UK Biobank. Each panel displays the mean and 95% CI of the PRS prediction R^2^ (y-axis) for Meta-PRS in each simulated scenario using either C+T or LDpred to generate the external PRS. The weights were obtained using linear regression (lm) or the square root of the training effective sample size (neff). In the case of the linear regression, the weights are trained in an independet validation dataset (see Table 1). The prediction R^2^ was transformed to the liability-scale using a population prevalence of 0.01 (ASD), 0.05 (ADHD), 0.15 (MDD UKB), 0.05 (T2D), 0.01 (AN), 0.03 (CAD), 0.01 (SCZ), 0.07 (BC), 0.01 (BD) and 0.08 (MDD iPSYCH). Mean and 95% CI of the AUC were obtained from 10k non-parametric bootstrap samples of the 5 cross-validation subsets.

